# HIV-1 latent infection triggers broader epigenomic and transcriptional changes in protein-coding and long non-coding RNAs than active infection of SupT1 cells

**DOI:** 10.1101/2022.08.10.503487

**Authors:** Gabrielle Lê-Bury, Yao Chen, Jordan M. Rhen, Jennifer K. Grenier, Amit Singhal, David G. Russell, Saikat Boliar

**Affiliations:** Department of Microbiology and Immunology, College of Veterinary Medicine, Cornell University, Ithaca, NY, USA; A*STAR Infectious Diseases Laboratories, Agency for Science, Technology and Research, Singapore, Singapore; Transcription Regulation and Expression Facility, Cornell University, Ithaca, NY, USA

**Keywords:** HIV-1, latency, epigenome, transcriptome, lncRNA

## Abstract

Latent HIV-1 infection poses a major challenge in complete viral remission and cure. HIV-1 latency is a multi-dimensional, dynamic process and many aspects of how the viral latency is established and maintained still remains incompletely characterized. Here, we have investigated the host chromatin organization and transcriptomic changes in active- and latently-infected SupT1 cells. We employed an *in vitro* model of HIV-1 latency in SupT1 cells using a dual-reporter virus, HIV_GKO_, which enables high purity sorting and characterization of active- and latently-infected cells. We found a significant divergence in chromatin organization and gene expression pattern between active and latent infection compared to uninfected cells. Latent infection results in a repressive reorganization of the host chromatin, while active infection leads to an overall increase in chromatin accessibility. A stronger correlation was also observed between chromatin accessibility and gene expression in latent infection, which was manifested in a greater alteration of the cellular transcriptome in latent than active infection, for both protein-coding and long-non-coding RNAs (lncRNAs). We identified a number of novel lncRNAs associated with either active and latent infection. A reversal in expression pattern of latency-associated lncRNAs following PMA-induced reactivation indicated their infection-state-specific expression and potential roles in HIV-1 latency. Taken together, this integrated, comparative study revealed that latent HIV-1 infection requires a substantially greater alteration in cellular epigenome and transcriptome. Understanding of the distinct cellular states conducive to active and latent infection may support devising strategies for specific modulation of host cellular functions as a curative intervention for HIV-1.

## INTRODUCTION

One of the major challenges in the cure for HIV-1 infection is the persistence of latently-infected cells. T-lymphocytes and macrophages can both serve as reservoirs for viral persistence, but resting memory CD4^+^T cells are considered the main cell type for HIV-1 latency (1–5). Current combination anti-retroviral therapy (cART) regimens can effectively impede viral replication and disease progression, but cannot eliminate the latently-infected cells. Even after prolong cART therapy, in the event of a treatment interruption or suspension, the virus rebounds rapidly from persistently-infected reservoir cells. The mechanisms of HIV-1 latency are not completely understood, but it is evident that transcriptional silencing of the provirus is multifactorial, regulated intrinsically by both the virus and the host cell (6, 7). Epigenetic repression of the proviral promoter region plays a role in viral transcriptional blockade (8–10). The site of viral integration into host genome is another determinant of viral gene expression and latency (11). Inadequacy of transcription factors such as NF-kB, NFAT and p-TEFb can impart viral latency as well (12, 13).

HIV-1 replication in infected cells relies on the host cellular machinery, therefore a comprehensive understanding of the cellular environment that supports establishment and maintenance of either viral replication or latency is crucial for effective designing of curative interventions for HIV-1. Multiple studies have undertaken epigenomic, transcriptomic and proteomic analysis of HIV-1 infected cells, both *in vivo* and *in vitro*, to interrogate the cellular and molecular changes that occur during active or latent viral infection (14–17). HIV-1 can modulate host defense pathways such as innate or adaptive immune response and immune activation at both epigenomic and transcriptomic levels (18, 19). Differential usage of cell metabolic pathways is also evidenced in active or latent infection (20).

Apart from protein-coding genes (PCGs), HIV-1 infection also modulates expression of various non-coding RNAs such as microRNAs and long non-coding RNAs (lncRNAs) (21, 22). In recent years, a number of lncRNAs have been identified with key roles in HIV-1 replication and pathogenesis. For example, a group of lncRNAs including *Malat1, HEAL, NRON* and *NEAT1* have been proven crucial in regulating HIV-1 replication and latency (23–26). On the other hand, lncRNAs *SAF* and *lincRNA-p21* can promote viral persistence by promoting survival of the infected cells (27, 28). Despite these multifaceted studies, a knowledge gap still exists. Recent technological advancements in high-throughput analysis and its usage in direct, unbiased comparison of active- and latently-infected cells relative to uninfected ones may help advance our understanding of the specific molecular pathways that shape the active and latent HIV-1 infection.

Absence of definite cellular markers to identify latently-infected cells as well as their rarity *in vivo* greatly hinder in-depth study of the viral reservoir cells obtained directly from HIV-1-infected individuals. Most of the advancements in our fundamental understanding of the mechanisms of HIV-1 latency have been achieved with *in vitro* models. A number of *in vitro* models of HIV-1 latency are currently in use and their merits and limitations have been discussed extensively (29). In this study, we used a single-round, dual-reporter HIV-1 virus, HIV_GKO_, which encompasses two distinct fluorescent readouts: one for viral infection and provirus integration (mKO2, under a constitutive cellular promoter EF1α) and another to indicate active viral transcription (codon-switched GFP under viral LTR promoter) (30). Infection with HIV_GKO_ virus thereby provides a distinct advantage of simultaneous identification of both actively-infected (viral transcription-active mKO2^+^GFP^+^) and latently-infected (viral transcription-inactive mKO2^+^GFP^-^) cells from within a pool of infected cells. We have utilized this *in vitro* model of HIV-1 infection in SupT1 cells and carried out an integrated, comparative analysis of the active- and latently-infected cells to that of virus non-exposed, uninfected cells to gain an insight into the disparate epigenomic and transcriptional landscape that distinguishes the two states of HIV-1 infection.

In this study, the ATAC-seq results revealed a drastic repression of host cell chromatin accessibility in latently-infected cells, while active HIV-1 infection enhanced overall accessibility. Consequently, there was also a greater degree of modulation of gene expression, as revealed by RNA-seq, in latent infection that affected biological pathways such as RNA metabolism, transcription and cell cycle. Through guilt-by-association analysis, we also identified several novel lncRNAs with potential roles in HIV-1 replication and latency. Reactivation of latent cells induced a trend-reversal in expression of latency-associated lncRNAs, indicating their infection-state-specific expression. Overall, our study revealed a greater impact of latent than active HIV-1 infection on chromatin organization and gene expression in T cells.

## RESULTS

### Marked repression of host chromatin organization in HIV-1 latent infection

HIV-1 latency in CD4^+^T cells is characterized by transcriptional silencing of viral genes, which is often attributed, at least in part, to limited host transcription factor accessibility to the proviral promoter region (31). To investigate whether such epigenomic alterations also extend to the host genome, we compared the overall host chromatin architecture in active- and latently-HIV-1-infected SupT1 cells to that of uninfected cells by ATAC-seq. Four days post-infection (dpi), HIV_GKO_-infected SupT1 cells were flow-sorted into the two populations: active (mKO2^+^GFP^+^) and latent (mKO2^+^GFP^-^) HIV-1-infected cells (Fig S1A). Uninfected, flow-sorted SupT1 cells that were also processed as controls (Fig S1B). Flow-sorted cells from three independent experiments were processed for ATAC-seq. After aligning the quality-filtered, paired-end reads to the human reference genome (GRCh38), we used irreproducible discovery rate (IDR) framework to identify highly reproducible peaks (IDR<0.05 between individual samples and pooled data of triplicates). This resulted in an average of 49000 peaks in each sample and a total of 131019 peaks after merging peaks across all samples. The distribution of chromatin accessibility peaks was comparable among the samples with most reads located near the transcription start sites (TSS) (Fig 1A). Principle Component Analysis (PCA) revealed separation of cells based on infection status, suggesting active and latent infection-specific chromatin organization.

**Figure 1:**
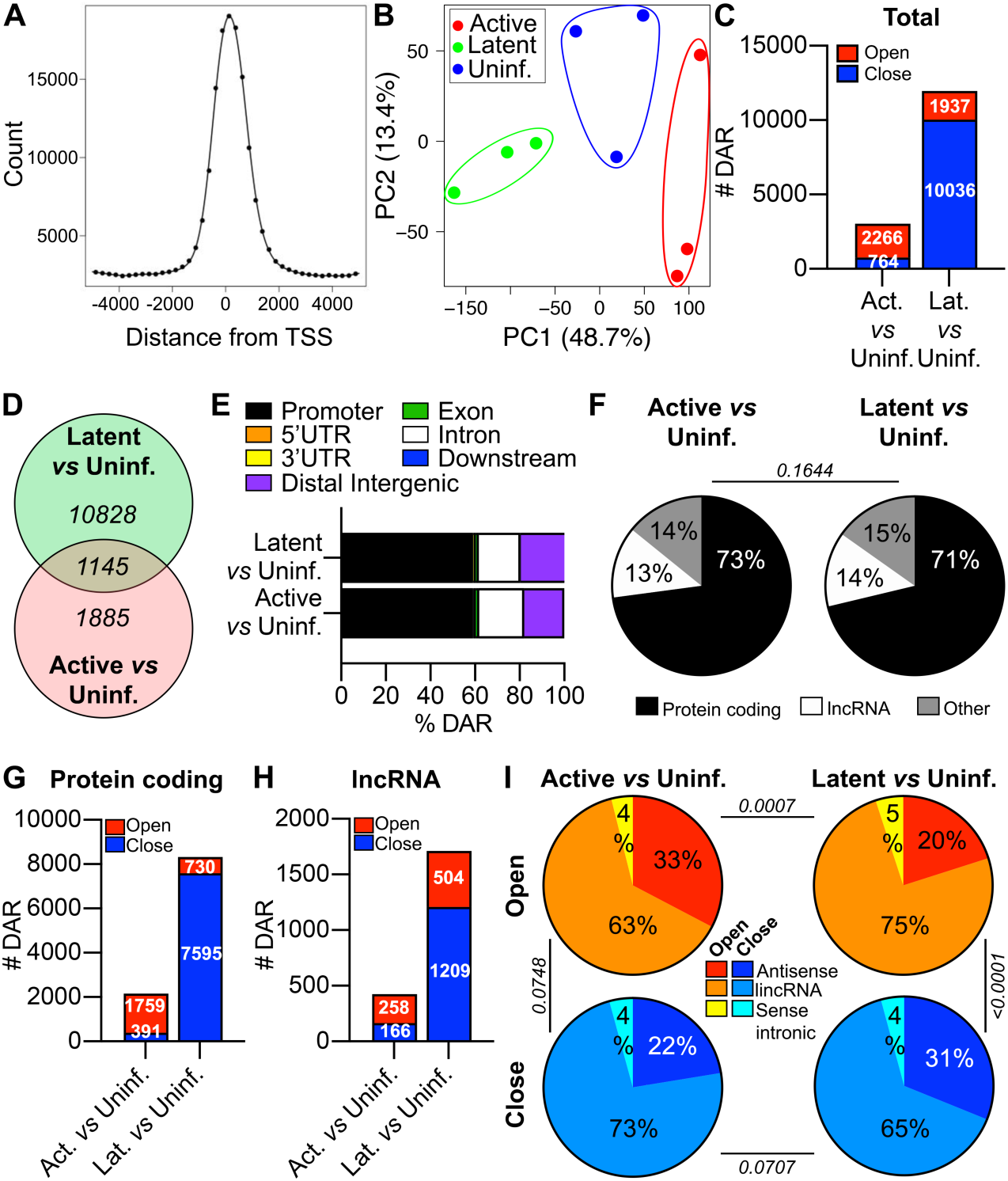
Chromatin accessibility in active and latent HIV-1-infection of SupT1 cells. (A) TSS enrichment plot of ATAC-seq peaks for combined datasets of active, latent and uninfected SupT1 cells. (B) PCA plot of ATAC-seq peaks from uninfected (blue), active-infected (red) and latent-infected (green) SupT1 cells. (C) Number of open (red) and close (blue) DARs (FDR < 0.05, fold change > 1.5) in active infection (left) or latent infection (right) compared to uninfected SupT1 cells. (D) Venn diagram showing common or exclusive DARs for active (red) or latent (green) infection compared to uninfected SupT1 cells. (E) Stacked bar plots indicating the genomic feature distributions of DARs in active (bottom) and latent (top) infection compared to uninfected SupT1 cells. Promoter (black), 5’UTR (orange), 3’UTR (yellow), distal intergenic (purple), exon (green), intron (white), downstream (blue) are color-coded. (F) Percentages of DARs associated with PCGs (black), lncRNAs (white) and other transcripts (grey) in active (left) and latent (right) infection compared to uninfected SupT1 cells. Chi-square test was used to compare distribution of frequencies. (G-H) Numbers of DARs (FDR < 0.05, fold change > 1.5) associated to PCGs (G) and lncRNA (H) that are open (red) or close (blue) in active (left) or latent (right) infection compared to uninfected SupT1 cells. (I) Percentages of lncRNA subclasses open (top) or close (bottom) in active (left) and latent (right) infection compared to uninfected SupT1 cells. Chi-square test was used to compare distribution of frequencies.

Comparison of chromatin accessibility profiles of the active- and latently-infected cells against the uninfected SupT1 cells revealed that latent HIV-1 infection led to significantly greater (∼4-fold) alterations in chromatin organization than in active HIV-1 infection, when compared to uninfected cells (Fig 1C). Overall, we identified 3030 and 11973 annotated regions of the genome in active- and latently-infected cells, respectively, that were significantly differentially accessible (DAR, Differential Accessible Regions, fold change > 1.5, FDR < 0.05) in comparison to uninfected cells (Fig 1C). Notably, in active HIV-1 infection, the proportion of open DARs was almost 3-fold higher than closed ones (Fig 1C). On the contrary, in latently-infected cells, the number of closed regions far exceeded (5-fold) the number of open DARs (Fig 1C). We noticed 1145 DARs that shared altered accessibility between active and latent infection in comparison to uninfected cells (Fig 1D). With regard to distribution, majority of DARs were associated with the promoter regions (TSS +/− 3kb) of genes, accounting for about 59% of the DARs in both active and latent infection (Fig 1E). Next, we classified the DARs based on their association to known annotated genes, either PCGs or lncRNAs. In both states of infection, impact on PCGs and lncRNAs remained comparable (Fig 1F). After segregating the DARs into either PCG- or lncRNA-associated DARs, the overall trend in epigenomic alteration was maintained; more open chromatin regions were observed in active infection whereas latent infection featured more closed chromatins for both PCGs and lncRNAs (Fig 1G-H). Within lncRNAs, the major affected subclasses were: antisense, intergenic (lincRNAs) and sense intronic lncRNAs (Fig 1I). The distribution of DARs associated to different lncRNA subclasses were comparable, except the proportion of open lincRNA-associated DARs was significantly overrepresented in latent infection (Fig 1I). Together, our data depicts an overall increased chromatin accessibility in active HIV-1 infection, whereas latent infection led to a collective decrease in chromatin accessibility. This indicates that HIV-1 latent infection in SupT1 cells triggered a more extensive, largely repressive reorganization of the host chromatin than in actively-HIV-1-infected cells.

### Broader transcriptional changes in HIV-1 latent infection

Since majority of DARs were in the promoter regions of either a PCG or lncRNA genes (Fig 1E), we investigated how the chromatin reorganization could affect the host transcription during the two states of HIV-1 infection. RNA-sequencing was performed with total cellular RNA extracted from the flow-sorted uninfected, active- and latently-infected SupT1 cells from two independent experiments. A total of 15724 RNA transcripts were found to be expressed across the samples, of which 11812 were PCGs and 2191 were lncRNAs (Fig 2A). Among the different subclasses, lincRNAs and antisense lncRNAs represented the two major classes, accounting for about 45% and 37%, respectively, of the overall expressed lncRNAs (Fig 2A). As expected, the median expression level of lncRNAs was more than 2-fold lower than that of the PCGs (Fig 2B-C).

**Figure 2:**
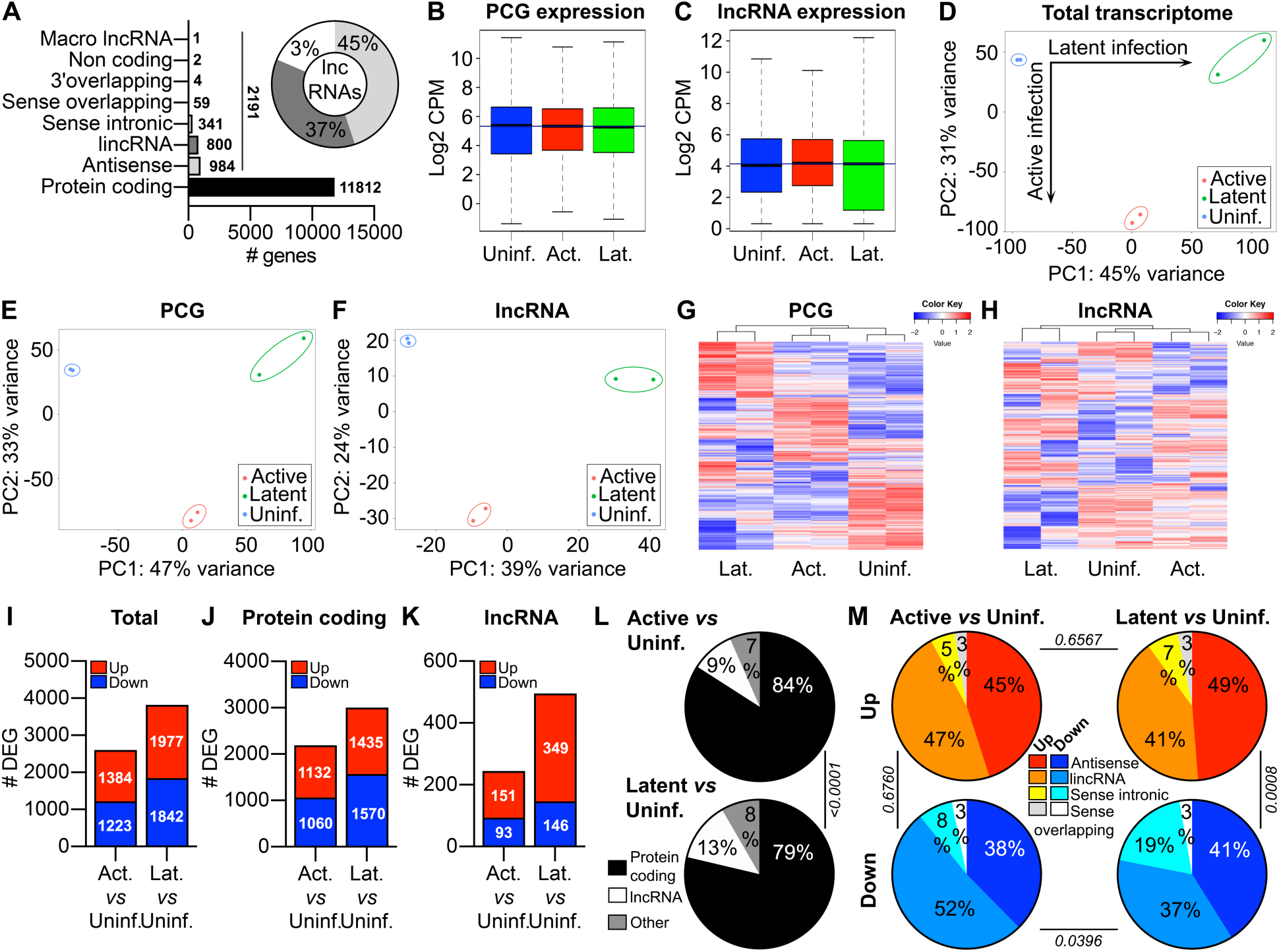
Expression of PCGs and lncRNAs in active and latent HIV-1 infection of SupT1 cells. (A) Distribution of transcript biotypes identified (cpm > 1 in at least 50% of samples) in RNA-seq datasets. Diagram shows the proportions of lncRNA subclasses. (B-C) Bar plots showing expression [log2 (cpm)] of PCGs (B) and lncRNAs (C) in uninfected (blue), active-infected (red) and latent-infected (green) SupT1 cells. (D-F) PCA plots representing the total transcriptome (D), PCGs (E) and lncRNAs (F) expression profiles in uninfected (blue), active-infected (red) and latent-infected (green) SupT1 cells. (G-H) Heatmaps of all detected PCGs (G) and lncRNAs (H) expression. (I-K) Numbers of DEGs (FDR < 0.05, fold change > 2) for total transcripts (I), PCGs (J) and lncRNAs (K) that are up-regulated (red) or down-regulated (blue) in active (left) and latent (right) infection compared to uninfected SupT1 cells. (L) Percentages of PCGs (black), lncRNAs (white) and other transcripts (grey) that are differentially expressed in active (left) and latent (right) infection compared to uninfected SupT1 cells. Chi-square test was used to compare distribution of frequencies. (M) Percentages of lncRNA subclasses that are up-regulated (top) or down-regulated (bottom) in active (left) and latent (right) infection compared to uninfected SupT1 cells. Chi-square test was used to compare distribution of frequencies.

To determine how the state of HIV-1 infection affected the global transcriptome of SupT1 cells, we performed unsupervised analysis among the uninfected, active- and latently-infected cells. PCA of the total transcriptome showed a robust segregation amongst the three different cell populations (Fig 2D). Next, we separated the PCGs and lncRNAs and analyzed their expression profiles singularly either by PCA or unbiased hierarchical clustering to understand their relative contributions to the divergent host response to HIV-1 infection state (Fig 2E-H). These analyses revealed that the distinction between uninfected and active- or latently-infected cells was accurately reflected in expression of both lncRNAs and PCGs. Interestingly, for both PCGs and lncRNAs, latently-infected cells clustered on a distinct branch on the dendrogram, depicting a higher degree of transcriptomic reprogramming in latent- than actively-infected cells in comparison to uninfected ones (Fig 2G-H). These expression profiles indicate that active and latent HIV-1 infection, akin to PCGs, imparts a unique lncRNA expression signature that effectively distinguishes the state of viral infection in SupT1 cells.

To measure the magnitude of host transcriptional changes that occur during HIV-1 infection of SupT1 cells, we identified the RNA transcripts that were differentially expressed (DE, fold change > 2 and FDR < 0.05) in active- and latently-infected cells relative to uninfected ones. A total of 2607 genes were DE during active HIV-1 infection compared to 3819 genes in latent, indicating broader host transcriptomic change in HIV-1 latency than active infection (Fig 2I). The pattern was similarly reflected in both DE PCGs and lncRNAs, although the difference was more marked for lncRNAs with twice as many DE lncRNAs in latently-infected cells than in active infection (Fig 2J-K). Furthermore, DE lncRNAs accounted for a larger portion (13%) of total dysregulated genes in latently-infected cells than in active (9%) infection (Fig 2L). Comparing the directionality, the number of up- or down-regulated PCGs were proportionate within each group of virus-infected cells (Fig 2J). However, a considerably higher number of lncRNAs were up-regulated than down-regulated in both states of infection (active: 1.6-fold, latent: 2.4-fold; Fig 2K). The distributions of different classes of DE lncRNAs were mostly similar between active and latent infections, except a significant higher proportion of sense intronic lncRNAs were down-regulated in latent infection (Fig 2M). Overall, these data show a greater dynamic shift in the host transcriptome during latent infection than in active HIV-1 infection, particularly in lncRNA expression.

### Superior correlation between chromatin accessibility and transcriptomic changes in HIV-1 latent infection

To understand the relationship between changes in chromatin accessibility and gene expression in active- and latently-HIV-1-infected SupT1 cells, we first estimated the number of genes that were mutually enriched in both DARs in ATAC-seq and DEGs in RNA-seq. Association of gene expression to changes in a corresponding genomic region chromatin accessibility showed a disparate pattern based on gene biotype as well as state of HIV-1 infection. Irrespective of infection status, a significantly larger fraction of differentially expressed DE PCGs (active: 15%, latent 60%) also had an associated DAR than that of lncRNAs (active: 4% latent: 18%), indicating a larger impact of chromatin reorganization on expression of PCGs than on lncRNAs (*p-value* < 0.0001, Fig 3A). However, when compared between active- and latently-infected cells, the proportion of DEGs (for both PCGs and lncRNAs) with an associated DAR, was significantly higher in latent infection (PCGs: 60%, lncRNAs: 18%) than in active HIV-1 infection (PCGs: 15%, lncRNAs: 4%) of SupT1 cells (*p-value* < 0.0001, Fig 3A). This suggests a greater role of chromatin reorganization in regulation of gene expression in latently-infected cells. Furthermore, when we compared the magnitude of changes (log2FC) in DARs and DEGs of the mutually altered transcripts, we observed a weak but significant correlation only in latently-infected cells, for both PCGs and lncRNAs (Fig 3B-E). These data suggest to a possible inherent difference in chromatin-mediated regulation of gene expression in active- and latently-infected cells, wherein a greater degree of correlation exists between chromatin remodeling and gene expression in latent HIV-1 infection.

**Figure 3:**
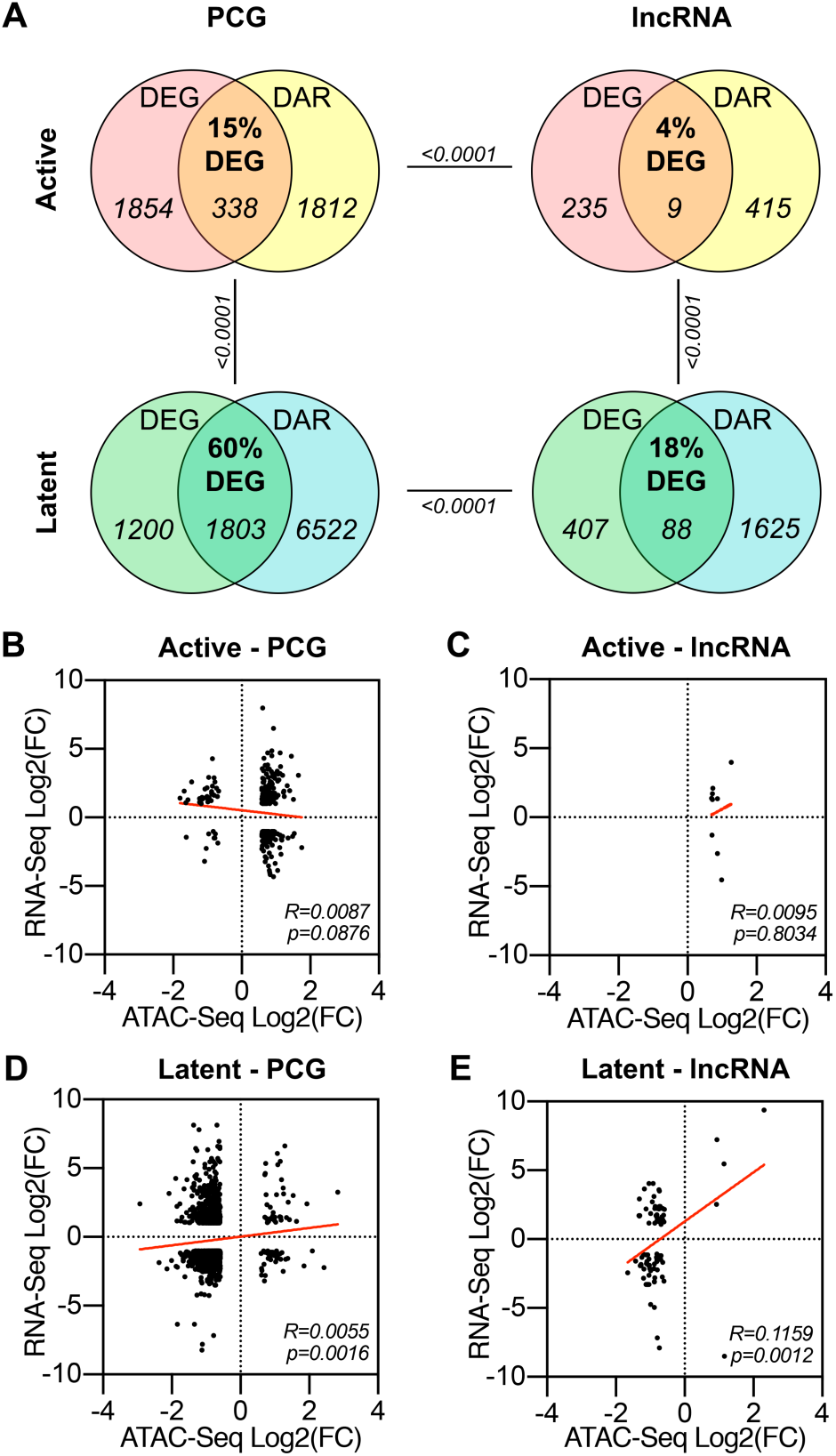
Correlation between chromatin accessibility (ATAC-seq) and expression (RNA-seq) of PCGs and lncRNAs. (A) Venn diagrams of significant DEGs and DARs, corresponding to PCGs (left) and lncRNAs (right) in active (top) and latent (bottom) infection compared to uninfected SupT1 cells. (B-E) Correlations of mutually altered DARs and DEGs associated to PCGs (B, D) and lncRNAs (C, E) in active (B, C) and latent (D, E) infection compared to uninfected SupT1 cells. Linear regression analysis was performed to obtain the R-squared coefficient of the slope (*R*) and *p-value* (*p*).

### Dysregulation of RNA metabolism, transcription and cell cycle pathways in HIV-1 infection

We next investigated the biological pathways that were specifically impacted in active- and latent-infected SupT1 cells. We first sought to identify the PCGs and lncRNAs that showed unique expression in either active or latent HIV-1 infection and excluded the ones that were mutually altered in both HIV-1 infection states. We analyzed the DE gene-sets with a proportional Venn diagram to identify the unique and shared transcripts among the cell populations (Fig 4A-B). RNA transcripts that were exclusively DE in active infection *versus* uninfected cells, as well as the genes that were mutually up- or down-regulated in active infection compared to both uninfected and latent-infected cells were considered to be active infection-specific (AI-PCG or AI-lncRNA), as depicted in Fig 4A-B. Similarly, DE genes that were unique to latently-infected cells compared to either active or uninfected cells were designated as latent infection-specific (LI-PCG or LI-lncRNA) (Fig 4B). Based on these criteria, we identified a total of 1008 PCGs and 104 lncRNAs that were specific to active HIV-1 infection. A larger pool of 1869 PCGs and 371 lncRNAs were found to be exclusively altered in latent HIV-1 infection. A hierarchical heatmap analysis of the identified active- and latent-infection-associated PCGs and lncRNAs clearly shows their expression specific to their respective state of viral infection (Fig 4C-D).

**Figure 4:**
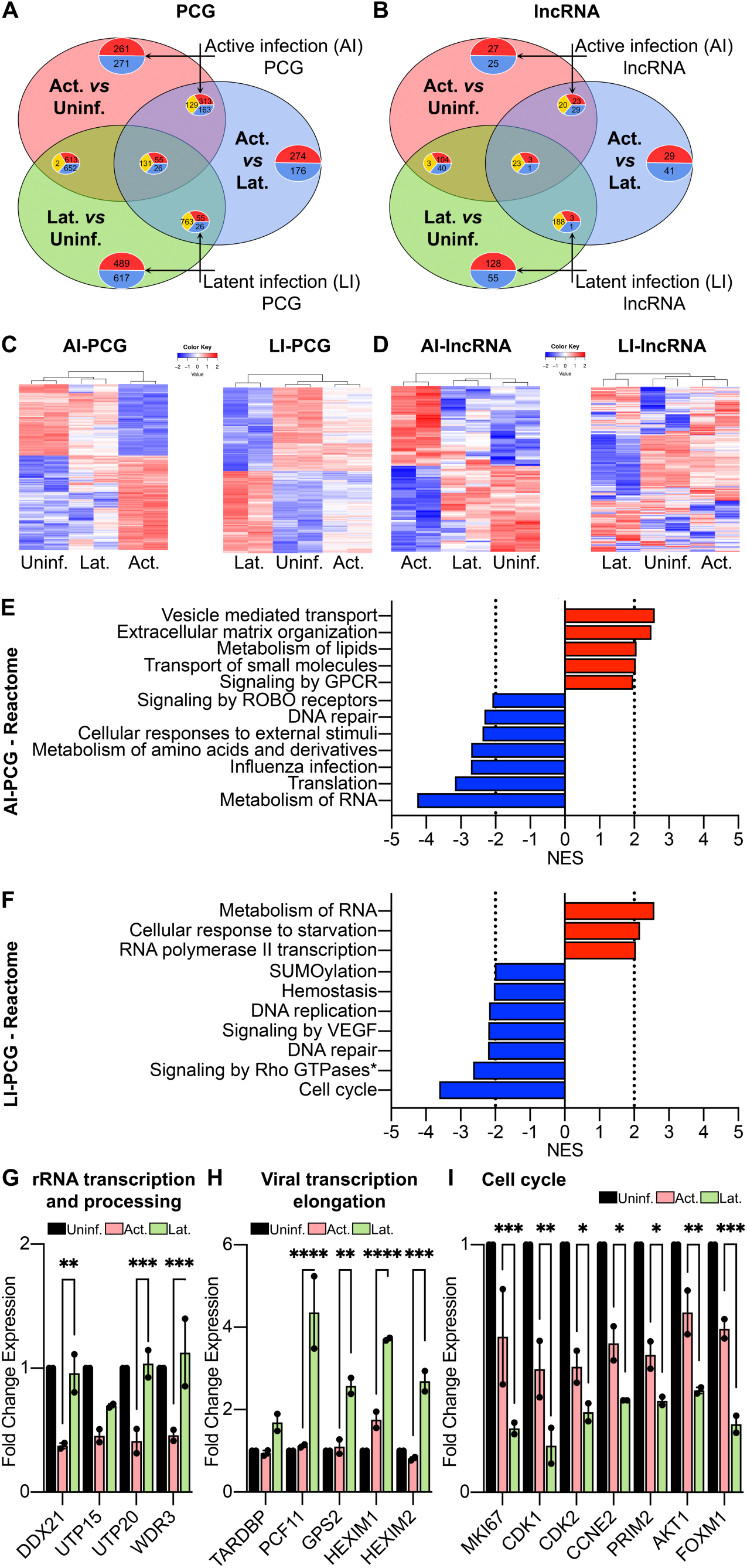
Biological pathways enriched in active and latent HIV-1 infection of SupT1 cells. (A-B) Venn diagrams of DEGs (FDR < 0.05, fold change > 2) amongst “active *versus* uninfected” (light red), “latent *versus* uninfected” (light green) and “active *versus* latent-infected” (light blue) SupT1 cells for PCGs (A) and lncRNAs (B). Numbers of DEGs that are up-regulated (red), down-regulated (blue) or contra-regulated (yellow) are indicated within plots. (C-D) Heatmaps of expression of PCGs (C) and lncRNAs (D) specific to active infection (AI-PCG or AI-lncRNA on the left) and latent infection (LI-PCG or LI-lncRNA on the right). (E-F) Biological pathways enriched (NES > 2) within the Reactome database for DE-PCGs (FDR < 0.05, fold change > 2) specific to AI-PCG (E) and LI-PCG (F) (* Signaling by Rho GTPases, Miro GTPases and RHOBTB3). (G-I) Relative fold change in expression of genes involved in rRNA Transcription and processing (G), viral transcription and elongation (H) and cell cycle (I) for active (red) and latent (green) infection compared to uninfected (black) SupT1 cells. Two-way ANOVA Fisher’s LSD test was used to generate p-value (*=p<0.03; **=p<0.002; ***=p<0.0002; ****=p<0.0001).

To understand how biological processes are impacted in active and latently-HIV-1 infected SupT1 cells, we performed pre-ranked gene set enrichment analysis (GSEA) with the active and latent infection-associated PCGs using the Reactome database of canonical pathways. Pathways with a normalized enrichment score > 2 and p-value < 0.01 were considered significantly altered (Fig 4E-F). Cellular transport of small molecules as well as vesicle-enclosed cargo were markedly increased in actively-infected cells. Active viral replication also up-regulated G-protein coupled receptors (GPCR)-mediated signal transduction in those HIV-1-infected SupT1 cells (Fig 4E). On the other hand, pathways associated with DNA repair, responses to external stimuli and metabolism of amino acid derivatives were down-regulated in actively-infected cells (Fig 4D). Interestingly, biological pathways associated with metabolism of RNAs showed opposing regulatory signatures in active and latent HIV-1-infection (Fig 4E-F). In latently-infected SupT1 cells, RNA metabolic processes along with RNA polymerase II-mediated transcription were up-regulated, whereas DNA damage response, DNA replication and cell cycle pathways were significantly down-regulated (Fig 4F).

#### RNA metabolic processes are suppressed in active infection, but up-regulated in latent infection

Metabolism of RNA is a super-pathway that encompasses important biological processes including conversion of nascent RNA transcripts into mature mRNAs (capping, splicing, polyadenylation) as well as editing and decaying of mRNAs. In active HIV-1 infection, ribosomal RNA processing was the predominant subnetwork that was significantly down-regulated, as majority (36/70 genes) of the RNA metabolism-enriched genes were involved in this process. Of note, HIV-1 infection of T-lymphocytes impairs ribosome biogenesis by inhibiting pre-rRNA processing (15). Genes related to rRNA transcription and processing, such as *DDX21, UTP15, UTP20, WDR3* were down-regulated in actively-infected SupT1 cells (Fig 4G). Consequent to the inhibition of ribosomal biogenesis, active viral replication also led to a suppression of the host translational machinery (Fig 4E). On the other hand, in latently-infected cells, RNA polymerase II-dependent transcription and metabolic processing of mRNAs was elevated compared to uninfected cells (Fig 4F). Intriguingly, it is known that one of the main contributing factors in HIV-1 latency is suppression of viral transcription. This indicates that inhibition of HIV-1 viral transcription in latently-infected cells is independent of overall suppression of cellular transcription. Indeed, expression of several transcription factors that enhances cellular transcription such as zinc finger proteins (*ZNF470, ZNF573*) and mediator complex subunits (*MED6, MED26, MED31*) was increased in latently-infected SupT1 cells (data not shown). On the other hand, suppressors of viral transcription such as *TARDBP*, which inhibits viral transcription by binding to the HIV-1 TAR element (32), was increased (1.7-fold) in latent infection (Fig 4H). Furthermore, repression of HIV-1 gene expression is often attributed to promoter-proximal stalling of RNA polymerase II and premature termination of viral transcription by increased recruitment of cleavage and polyadenylation factors to the viral transcription start site. Expression of several factors implicated in inhibition of viral transcription elongation, such as *PCF11* (4.3-fold), *GPS2* (2.6-fold), *HEXIM1* (3.7-fold) and *HEXIM2* (2.7-fold) were significantly up-regulated in latently-infected SupT1 cells compared to actively-infected cells (Fig 4H). Our results demonstrate that interference of HIV-1 transcription in latently-infected SupT1 cells is virus-specific and not obligated to an overall down-regulation in cellular transcription or post-transcriptional processes.

#### DNA replication and cell cycle pathways are markedly down-regulated in latent infection

Biological pathway analysis also showed a significant decrease in cell proliferation pathways in latently-infected SupT1 cells. Within the cell cycle, some of the differentially enriched pathways included cell cycle checkpoints, resolution of sister chromatid cohesion and mitotic spindle checkpoints. The cell proliferation marker *MKI67* as well as genes involved in G1-S and G2-M transition such as *CDK1, CDK2, CCNE2, PRIM2, AKT1, FOXM1* were significantly down-regulated in the latently-infected compared to actively-infected SupT1 cells (Fig 4I). HIV-1 accessory protein, Vpr, is known to impede cell cycle (33, 34) and accordingly, we observed a trend of lower expression of the cell cycle genes in actively-infected cells compared to uninfected ones (Fig 4I). However, the reductions were significantly more severe in latently-infected cells (Fig 4I), indicating a possible Vpr-independent cell cycle arrest in latent HIV-1 infection.

### Identification of putative associations of lncRNAs to cellular pathways dysregulated in active and latent HIV-1 infections

To gain an insight into the biological functions of the DE lncRNAs, most of which still remain functionally unannotated, we used a guilt-by-association analysis and constructed co-expression networks of the PCGs and lncRNAs. We analyzed all detectable RNA transcripts in the dataset by unsupervised Weighted Gene Co-expression Network Analysis (WGCNA) for hierarchical clustering using the “unsigned” mode to account for the opposing regulations that is frequently observed between interacting lncRNAs and PCGs. A total of 11 modules of significantly correlated transcripts were detected (Fig 5A). PCGs and lncRNAs did not cluster separately, rather remained coalesced within each module (Fig 5A-C). However, active- and latent-infection-associated genes remained mostly segregated into different modules, further supporting the transcriptional divergence in these two states of HIV-1 infection (Fig 5B-C). Modules 1, 2, 4, 8 and 9 included latency-associated genes, although vast majority of the latency-specific PCGs and lncRNAs were contained in modules 1 and 9 (Fig 5A-C). On the other hand, a substantial fraction of active infection-specific genes got assigned to module 3 and 7 (Fig 5A-C). Four modules (5, 6, 10 and 11) were populated with a mix of genes linked to both active and latent HIV-1 infection. We therefore focused our subsequent analysis on four major modules that encompassed the bulk of either active infection (Modules 3 and 7) or latent infection (Modules 1 and 9) associated genes.

**Figure 5:**
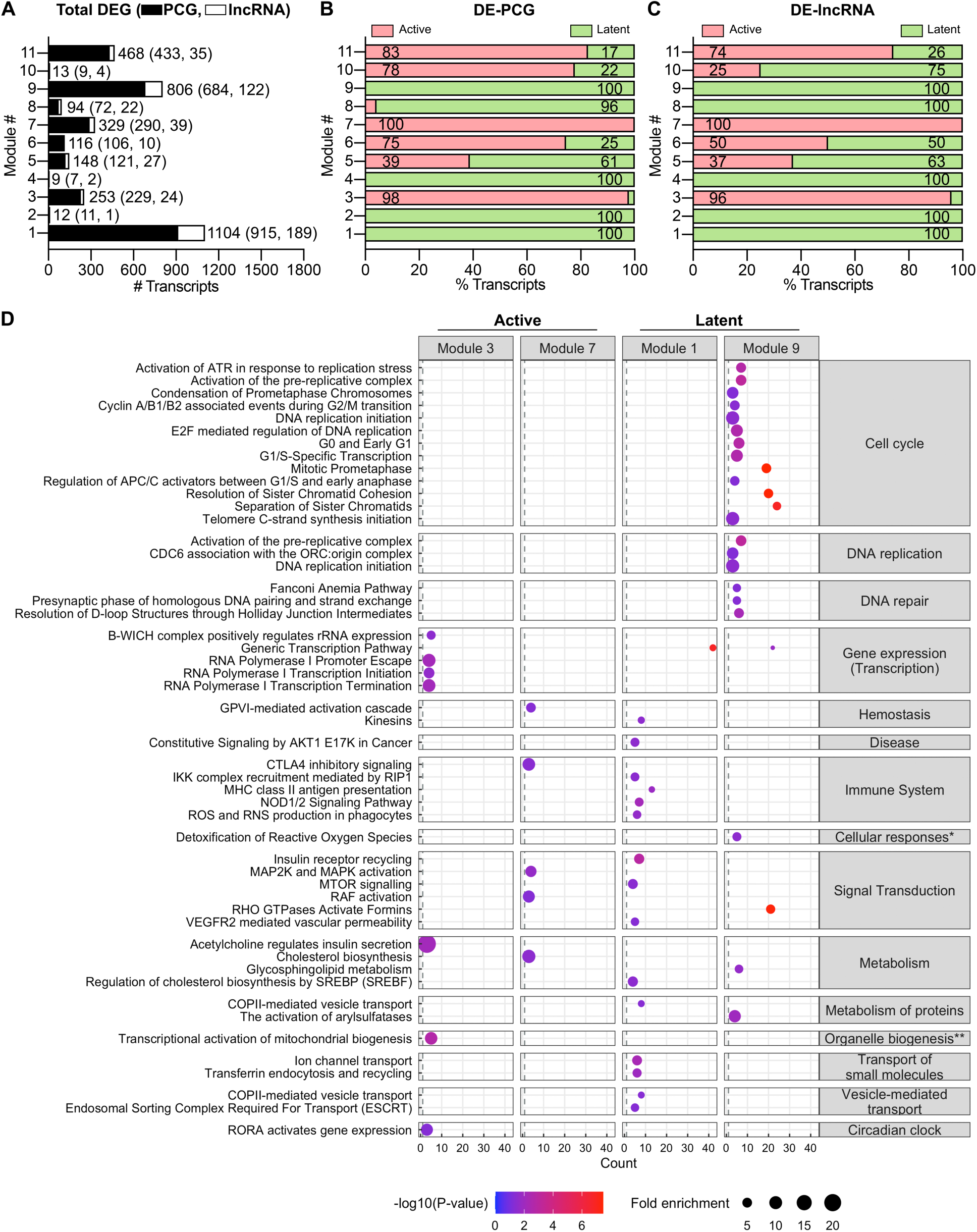
Identification of putative functions of DE-lncRNAs in active and latent HIV-1 infection of SupT1 cells. (A) Number of transcripts in each module identified by WGCNA co-expression analysis for PCGs (black) and lncRNAs (white). (B-C) Percentages of DE-PCGs (B) and DE-lncRNAs (C) in active (red) and latent (green) infection within each module. (C) Significantly enriched (*p-value* < 0.05) pathways within the Reactome database for active infection-specific modules 3 and 7; and in latent infection-specific modules 1 and 9. The dot size and color represent fold enrichment of the pathways and its significance (-log(*p*-value)), respectively. (*Cellular responses to external stimuli, **Organelle biogenesis and maintenance)

A mutual expression pattern between PCGs and lncRNAs often suggests shared biological functions and/or inter-molecular regulatory relations. To understand the potential functions of the lncRNAs, we analyzed the member PCGs within the active or latent infection associated modules for enrichment in canonical pathways based on the Reactome database (Fig 5D). Module 3, which contained predominantly active infection-specific genes, was enriched in annotations related to gene transcription pathways such as RNA polymerase I-mediated transcription initiation and termination as well as transcription of rRNAs and mitochondrial genes (Fig 5D). This suggests that the lncRNAs in module 3 could possibly be involved in RNA metabolism and translation, two processes that were found to be significantly down-regulated in actively-infected cells (Fig 4E). The active-infection-associated module 7 showed enrichment for pathways linked to immune responses, signal transduction and cholesterol metabolism. Metabolism of lipids was one of the pathways that were found to be up-regulated in actively-infected cells (Fig 4E). On the other hand, in latent infection-associated modules, a large number of PCGs was associated with pathways of gene transcription in module 1 and cell cycle, DNA replication or repair in module 9 (Fig 5D). Notably, gene transcription was up-regulated and cell cycle, DNA replication and DNA damage pathways were found to be down-regulated in latently-infected cells (Fig 4F).

Since RNA transcription/metabolism and DNA replication or cell cycle were the two thematic biological processes that were markedly modulated in active and latent HIV-1 infection of SupT1 cells, we sought to identify lncRNAs that were potentially associated with these pathways. For that, we identified the lncRNAs that showed the highest correlations (Pearson correlation coefficient > 0.99 and *p-value* < 0.01) within their respective modules, to the transcription and cell cycle-associated PCGs. This identified 61 lncRNAs showing the highest correlations with PCGs involved in gene transcription (Fig S2A, S2C). Similarly, 40 lncRNAs were found to be associated with PCGs annotated to cell cycle pathways (Fig S2B, S2D), indicating potential roles of these lncRNAs in these cellular pathways. In fact, the lncRNAs NRAV has been reported to play roles in transcription regulation of interferon-stimulated genes, while DLEU2 binds to Hepatitis B virus HBx protein to modulate gene transcription (35, 36). On the other hand, linc00665 is known to regulate cell cycle through inhibition of CDKN1C expression (37). The lncRNA SNHG3 has also been shown to regulate cancer cell proliferation through modulation of CyclinD1 and CDK1 (38). These published reports corroborate the correlative associations of the lncRNAs identified in this study.

### Reversal in expression of latency-associated lncRNAs following reactivation of latently-infected cells

We further investigated how latency reversing agent (LRA)-mediated stimulation of latently-infected T cells would affect the expression of lncRNAs that were identified as latency-associated and showed correlation to either gene transcription or cell cycle pathways (Fig S2A-B). Flow-sorted, latently-infected SupT1 cells were either treated with a PKC agonist LRA (PMA, 25ng/ml) or left untreated. After 24 hrs, we observed spontaneous reactivation of viral transcription (GFP expression) in a considerable fraction (mean, 46%) of untreated latently-infected SupT1 cells. PMA stimulation led to a further and significant increase in the number of reactivated (mean, 63%) cells (Fig 6A-B). At 24 hrs post-stimulation, we harvested the PMA-treated or untreated cells, extracted total cellular RNA and analyzed expressions of 4 transcription-associated lncRNAs and 5 cell cycle-associated lncRNAs that were significantly over-expressed in latent HIV-1 infection (Fig S2A-B). We also analyzed expression of NEAT1, a lncRNA known to increase during active HIV-1 replication (26). As expected, expression of NEAT1 tended to down-regulate in latent infection in comparison to active infection and PMA stimulation reverted its expression in reactivated cells to a level comparable to actively-infected cells (Fig 6C). Changes in expression of transcription-associated lncRNAs were disparate. While expression of the lncRNAs, RP11-255C15.3 and CTD-3222D19.12, showed a down-regulation after PMA stimulation; expression of C21orf91-OT1 and NRAV were either unchanged or increased, respectively, compared to latent infection (Fig 6D). On the other hand, cell cycle related lncRNAs, which were significantly up-regulated in latent infection, showed considerable reductions in their expression following PMA stimulation, including a significant decrease in expression of the lncRNA RP11-1C8.5 (Fig 6E). These data show a pattern in lncRNAs expression that is specific to HIV-1 infection state and are readily modulated upon change in the state of infection.

**Figure 6:**
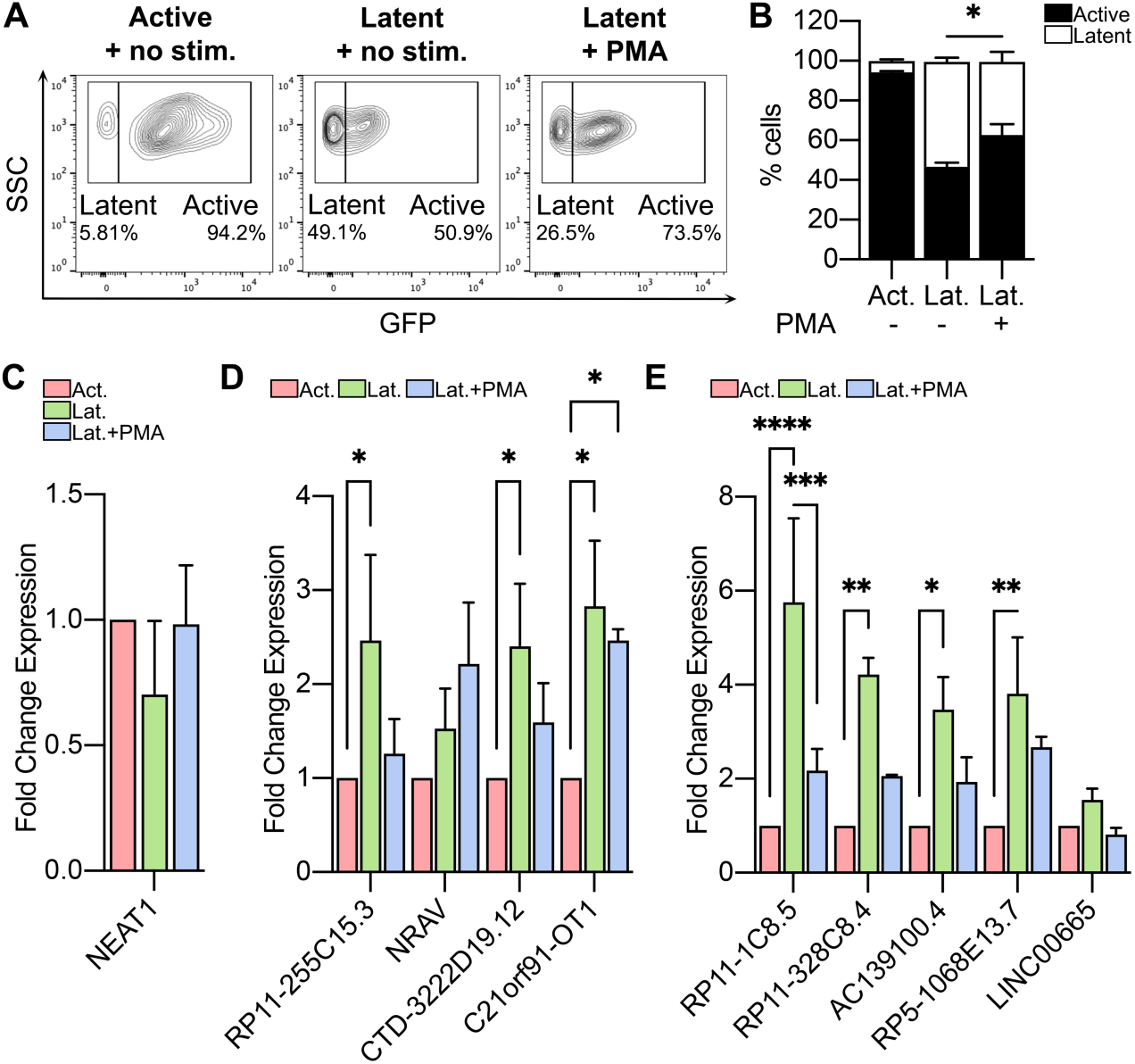
Altered expression of latency-associated lncRNAs following reactivation of latently-infected SupT1 cells. (A) Representative flow cytometry plots showing proportions of active and latent SupT1 cells in flow-sorted active cells without stimulation (left panel), latent cells without stimulation (middle panel) and latent cells with PMA (25ng/ml) stimulation for 24h (right panel). (B) Proportions of active and latent SupT1 cells from three independent experiments analyzed by two-way ANOVA test (*=p<0.03; **=p<0.002; ***=p<0.0002; ****=p<0.0001). (C-E) Relative fold changes in expression of *NEAT1* (C), lncRNAs associated to transcription pathway (D), and lncRNAs associated to cell cycle pathway (E) in unstimulated latent (green) and PMA-stimulated latent (blue) cells compared to actively-infected (red) SupT1 cells. Two-way ANOVA Fisher’s LSD test was used to compare lncRNA expressions from three independent experiments (\**=*p<0.03; **=p<0.002; ***=*p*-value<0.0002; ****=*p*-value<0.0001).

## DISCUSSION

HIV-1 infection is associated with a multitude of changes in host cellular functions that contribute to the establishment of either productive or latent infection. An in-depth understanding of such alterations is instrumental for crafting effective and precise therapeutics against HIV-1. The overall aim of this study was an integrative and comparative analysis of the epigenomic and transcriptomic changes that occur in active and latently-HIV-1-infected T cells. Several studies have previously investigated alterations in genomic, transcriptomic or proteomic profiles of HIV-1-infected T cells, often focused on either active- or latently-infected cells (16, 17, 39). However, in many instances, the analysis was confounded by use of a heterogenous mix of infected cells as the source material, without purification of active- and latently-infected cells. We used a well-established HIV-1 latency model of infection of SupT1 cells with a dual reporter virus, HIV_GKO_ that allows specific isolation and analysis of HIV-1 active- or latently-infected cells in utmost purity (30). It also facilitates investigation of cellular reprogramming at the early stages of active and latent infection without any extrinsic stimulations that are often necessary for HIV-1 infection of primary T cells. We employed a multi-omics approach encompassing both ATAC-seq and RNA-seq and directly compared flow-sorted active- or latently-infected SupT1 cells to uninfected ones for a more comprehensive insight. In congruence with previous findings, our results show that both epigenetic and transcriptomic profiles of HIV-1-infected T cells are quite distinctive to the state of viral infection and further expand the findings to demonstrate that the magnitude of changes is significantly greater in latently-infected than actively-infected SupT1 cells. We also demonstrated that the dynamic relationship between chromatin reorganization and gene expression was greater in latent HIV-1 infection.

The two most impacted cellular pathways that distinguished active and latent HIV-1 infection of SupT1 cells were gene transcription and cell cycle. HIV-1 infection, specifically the accessory protein, Vpr, is known to modulate cell cycle (33, 34). Here, we also observed a down-regulation of genes enriched in the cell cycle pathway in actively-infected SupT1 cells. However, the inhibition of gene expression and blockade in cell cycle pathway was significantly higher in latently-infected cells (Fig 4I). This suggests a possible Vpr-independent mechanism of cell cycle arrest in HIV-1 latent infection of T cells. The inhibition of cell cycle observed in latently-infected SupT1 cells is consistent with the fact that HIV-1 transcriptional silencing and latency in T-lymphocytes is facilitated by transition of virus-infected, activated CD4^+^T cells into resting, memory T cells (40). Furthermore, experimental induction cellular quiescence and restriction of cell cycle has also been shown to promote HIV-1 latency in T cells (41). However, the blockade in cell cycle pathway in latently-infected SupT1 cells observed in our study occurred in absence of immunological memory development or any external coercing factors, indicating an alternate regulatory pathway. Delineation of the exact mechanism of this restriction needs further investigation.

In this study, we comprehensively defined the expressed lncRNA repertoire in SupT1 cells. Almost 16% of all expressed RNA transcripts in SupT1 cells were annotated as lncRNAs (Fig 2A). We also investigated the effects of HIV-1 infection on lncRNA transcripts and identified putative functions of the differentially expressed lncRNAs in HIV-1 replication and latency. Indeed, integrated analysis of the epigenetic and transcriptomic profiles of lncRNAs in active- and latently-HIV-1-infected T cells had so far remained unexplored. Our study revealed a dramatic impact of HIV-1 infection on both the chromatin accessibility and expression of lncRNAs, which was more pronounced than PCGs. Divergent and distinct expression patterns of lncRNAs in active and latent infection provided evidence that, apart from PCGs, lncRNAs can also accurately reflect the state of HIV-1 infection in T cells. Moreover, we showed that LRA-mediated reactivation affects the dynamics of the lncRNAs that are differentially expressed during HIV-1 latency. We observed spontaneous reactivation in a considerable fraction of flow-sorted, latently-infected SupT1 cells. It is known that HIV-1 latently-infected cells remain in a pseudo-steady state, revolving between viral transcriptional silence and low level of viral reactivation. However, the higher level of spontaneous reactivation in the SupT1 cells observed in our study could be due to extraneous activation during the long flow-sorting period. Nonetheless, stimulation with PMA further increased the proportion of reactivated cells. Despite the spontaneous reactivation in untreated cells, expression of a number of lncRNAs that we tested showed a trend-reversal in the PMA-stimulated cells. Significant down-regulation in expression of the lncRNAs in PMA-reactivated cells further suggests a potential role of the lncRNAs in regulation of HIV-1 replication and/or latency. Future studies on the mechanism of functions of the lncRNAs identified here could provide additional modalities for manipulation HIV-1 infection and latency.

In summary, the multi-omics analysis in this study provides new insights into the distinct epigenomic and transcriptional states of active and latent HIV-1-infected cells and reveals a greater impact of HIV-1 latency than active infection on T cells. The knowledge can be leveraged in innovative modulations of cellular functions for therapeutic interventions of HIV-1.

## MATERIALS AND METHODS

### Cell lines and culture

HEK 293T, SupT1 cells were obtained from ATCC and TZM-bl cells from the NIH HIV Reagent Program. SupT1 cells were cultured in RPMI media supplemented with 10% fetal bovine serum (FBS) and 1% each of penicillin, streptomycin and glutamine. HEK 293T and TZM-bl cells were cultured in DMEM media with 10% FBS and 1% each of penicillin, streptomycin and glutamine.

### Virus production

HIV_GKO_ plasmid was obtained from Addgene (deposited by Dr. Eric Verdin). Viral stock was generated by co-transfecting HEK 293T cells with 10:1 ratio of HIV_GKO_ plasmid and a plasmid encoding VSV-G envelope using Lipofectamine 3000 reagent (Invitrogen). Culture supernatant was harvested 48hrs post-transfection, centrifuged (1500rpm, 5min) to remove debris and then filtered by passing through a 0.45μm pore-size polyvinylidene difluoride (PVDF) membrane. Viral stock was tittered by infecting TZM-bl cells and X-Gal staining for counting blue colony-forming units. Viral stocks were then aliquoted and stored in −80°C until use.

### SupT1 infection and sorting

SupT1 cells were infected with HIV_GKO_ virus at an multiplicity of infection of 0.1 by spinoculation (3000rpm, 30min, 30°C). Virus-infected or uninfected cells were cultured for 4 days in a 37°C incubator with 6% CO2. Uninfected or HIV-1 active-infected (mKO2^+^GFP^+^) and latent-infected (mKO2^+^GFP^-^) SupT1 cells were flow-sorted using a Biorad S3 cell sorter. Total RNA was extracted from flow-sorted cells using TRIzol reagent (Invitrogen) according to manufacturer’s protocol, followed by treatment with DNase (Turbo DNAse-free kit, Invitrogen).

### RNA-seq library preparation and sequencing

Ribosomal RNA (rRNA) was removed from the total RNA (50ng) using NEBNext rRNA Depletion kit v2 (Human/Mouse/Rat, New England Biolab), according to manufacturer’s instructions. The rRNA-depleted samples were used for stranded library preparation using NEBNext Ultra II Directional RNA Library Prep Kit for Illumina (New England Biolabs). The libraries were sequenced on a HiSeq2500 (Illumina) instrument with a targeted average output of 40M paired-end reads (150bp) per sample.

### RNA-seq data analysis

Raw RNA-seq reads were first analyzed with FastQC for quality control and the processed through Trim Galore for removal of low quality reads and the Illumina sequencing adapters (42). The trimmed, high-quality reads were aligned to the Ensembl human reference genome (GRCh38 v86) using the STAR aligner software (43). Raw reads count for each gene were obtained using −quantmode of GeneCounts. Aligned reads were annotated in accordance to ENSEMBL database and categorization of an RNA transcript as lncRNA was based on the ENSEMBL/GENCODE biotype classifications. RNA transcripts with counts per million mapped-reads (cpm) greater than 1 in at least 50% of the samples were considered detectable in the datasets. Differential expression analysis of the gene counts was carried out using the standard EdgeR pipeline in R (44). False discovery rate (FDR) < 0.05 and fold change (FC) >2 was considered significant. For pathways analysis, gene lists were pre-ranked based on their fold changes and *p*-values [sign(log2FC)*-log10(*p*-value)] prior to analysis by GSEA using the Reactome database of canonical pathways. Weighted gene co-expression network analysis (WGCNA) was performed using the “unsigned” mode following previously published method (45).

### ATAC-seq library preparation and sequencing

Uninfected or HIV-1 active-infected (mKO2^+^GFP^+^) and latent-infected (mKO2^+^GFP^-^) SupT1 cells were flow-sorted using a BioRad S3 sorter, as described above. ATAC-seq libraries were prepared from the flow-sorted cells following published protocols (46–48). Briefly, 100,000 cells were used to for transposition reaction (Illumina Tagment DNA TDE1 Enzyme and Buffer Kit). Tagmented DNA was purified with MinElute purification kit (Qiagen) and then used for subsequent library preparation by PCR amplification using unique dual-indexing (Illumina Nextera i5 common adapter and i7 index adapter) primers. ATAC-seq libraries were purified using AMPure SPRI beads (Beckman Coulter) and then sequenced on a NextSeq 2000 (Illumina) to a targeted, average depth of 50 million, 100bp paired-end reads per sample.

### ATAC-seq data analysis

Raw ATAC-seq reads were first processed by FastQC for quality control and then Illumina sequencing adapters were trimmed from paired-end reads using Cutadapt (42, 49). The trimmed reads were aligned to human reference genome GRCh38 using BWA-MEM and the resulting alignment output were analyzed by SAMStat for quality control and then sorted by SAMtools (50–52). Post-alignment filtering was performed by SAMtools, ‘MarkDuplicates’ program of Picard tools as well as ENCODE ATAC-seq script assign_multimappers.py to remove PCR duplicates, secondary alignments and unmapped reads. Quality control of fragment size distribution was carried out by ATACseqQC (53). To correct for the insertion of two adapters as Tn5 transposase binds as a dimer at the Tn5 tagmentation step, reads aligned to the positive strand were shifted 4bp downstream and those aligned to negative strand were shifted 5bp upstream. The aligned, filtered and processed reads were used for all subsequent analysis. Next, identification of chromatin accessible peaks was performed by MACS2, which was executed by ENCODE script encode_macs2_atac.py (smooth_win = 150, pval_thresh = 0.01, gensz = 3.21e9) (54). In order to carry out the irreproducible discovery rate (IDR) filtering in subsequent steps, we generated pooled FASTQ files for each group (actively-infected, latently-infected and uninfected) and performed read alignment, sorting, post-alignment filtering, Tn5 shifting as well as peak calling for the pooled data through the same pipeline as described above. For each group, IDR framework was applied to select highly reproducible peaks (IDR < 0.05) between individual replicates and the pooled data of the respective group. The selected peaks from all replicates of a group were then merged by BEDTools as the final set of IDR peaks for the group. Principal component analysis (PCA) was performed on peaks with variance among samples (IQR filtering was applied) by using ‘PCA’ function of FactoMineR. All the peaks were annotated to the nearest genes based on ENSEMBL annotation package ‘EnsDb.Hsapiens.v86’ by ChIPseeker (55). To perform differential peak analysis, we merged IDR peaks from all the groups and counted the reads that overlap merged peaks for each sample. Differential accessible regions between groups were then determined using EdgeR pipeline in R (44). FDR < 0.05 and FC> 1.5 was considered significant.

### Quantitative RT-PCR

For relative quantification, about 25 ng of DNA-free total RNA for each lncRNA was reverse-transcribed using iTaq cDNA Synthesis Kit (Bio-Rad) and then analyzed with ABI 7500 Fast Real-time PCR System (Applied Biosystem) using iTaq Universal SYBR Green Supermix (Bio-Rad). Expression of lncRNAs was normalized based on 3 house-keeping gene: GAPDH, 18S rRNA and U6. Primer sequences used in the assays are available in Supplementary Table 1.

### Statistical analysis

Gene expression between groups (normalized log counts from RNA-seq or fold change from qRT-PCR) were compared with a two-way ANOVA Fisher’s LSD test. A *p-value* less than 0.05 was considered significant. Distribution of frequencies of lncRNA subclasses between groups were compared with Chi-square test. GraphPad Prism software was used for the statistical analysis.

## Data availability

The RNA-seq and ATAC-seq datasets generated and used in this article have been deposited in and are accessible from GEO: GSE####

## Author contributions

GLB: experimentation, data analysis, manuscript – review and editing; YC: data analysis, manuscript – review and editing; JMR: experimentation; JKG: experimentation, consultation; AS: data analysis, manuscript – review and editing; DGR: manuscript – review and editing, supervision, funding acquisition; SB: conceptualization, experimentation, data analysis, manuscript – writing, supervision, funding acquisition.

## Acknowledgment

This work was supported by A*STAR ID Labs grant to AS and by National Institutes of Health grants R33 AI136097 to DGR and R21 AI157759 to SB.

