## Supplementary material for "HIV-1 latent infection triggers broader epigenomic and transcriptional changes in protein-coding and long non-coding RNAs than active infection of SupT1 cells": Fig S1

#### Supplementary Figure Legends

**Figure S1: Flow cytometry sorting of active- and latent-HIV-1-infected SupT1 cells.** (A-B) Representative flow cytometry plots showing the gating strategy for flow sorting the different populations of cells: uninfected, active and latent in $HIV_{GKO}$-infected SupT1 cells (A) and uninfected SupT1 cells (B).

**Figure S2: Transcription and cell cycle associated lncRNAs.**
A-B) Heatmaps of expression of lncRNAs associated with transcription pathway (A) and cell cycle pathway (B). * = lncRNAs selected for reactivation study.
(C-D) Correlation networks using Cytoscape for PCGs (white) and lncRNAs (up-regulated in red; down-regulated in blue) associated to transcription pathway (C) and cell cycle pathway (D). Red and green lines indicate inverse and positive correlations, respectively. Pearson correlation coefficient > 0.99 and *p*-value < 0.01.

#### Supplementary Table Legends

**Table 1: Primers sequences used for quantitative RT-PCR.**

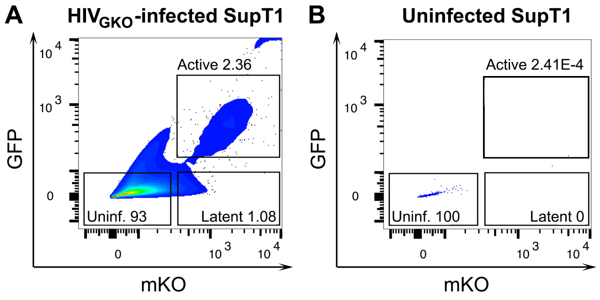
A
HIVGKO-infected SupT1
Active 2.36
Uninf. 93
Latent 1.08
GFP
mKO
B
Uninfected SupT1
Active 2.41E-4
Uninf. 100
Latent 0
GFP
mKO

### Figure S1

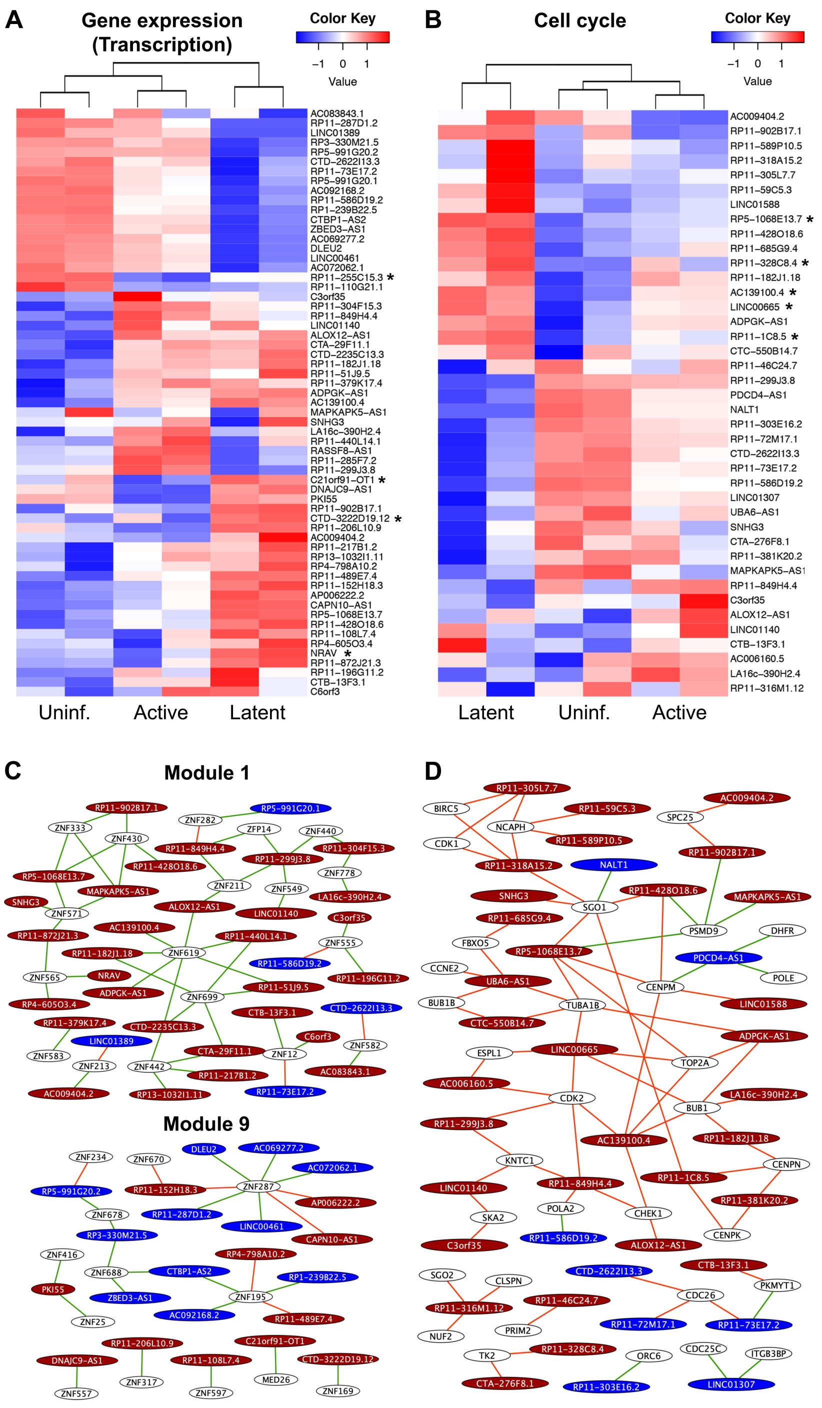
A
Gene expression (Transcription)
Color Key
−1 0 1
Value
AC083843.1
RP11−287D1.2
LINC01389
RP3−330M21.5
RP5−991G20.2
CTD−2622I13.3
RP11−73E17.2
RP5−991G20.1
AC092168.2
RP11−586D19.2
RP1−239B22.5
CTBP1−AS2
ZBED3−AS1
AC069277.2
DLEU2
LINC00461
AC072062.1
RP11−255C15.3 *
RP11−110G21.1
C3orf35
RP11−304F15.3
RP11−849H4.4
LINC01140
ALOX12−AS1
CTA−29F11.1
CTD−2235C13.3
RP11−182J1.18
RP11−51J9.5
RP11−379K17.4
ADPGK−AS1
AC139100.4
MAPKAPK5−AS1
SNHG3
LA16c−390H2.4
RP11−440L14.1
RASSF8−AS1
RP11−285F7.2
RP11−299J3.8
C21orf91−OT1 *
DNAJC9−AS1
PKI55
RP11−902B17.1
CTD−3222D19.12 *
RP11−206L10.9
AC009404.2
RP11−217B1.2
RP13−1032I1.11
RP4−798A10.2
RP11−489E7.4
RP11−152H18.3
AP006222.2
CAPN10−AS1
RP5−1068E13.7
RP11−428O18.6
RP11−108L7.4
RP4−605O3.4
NRAV *
RP11−872J21.3
RP11−196G11.2
CTB−13F3.1
C6orf3
Uninf.
Active
Latent
B
Cell cycle
Color Key
−1 0 1
Value
AC009404.2
RP11−902B17.1
RP11−589P10.5
RP11−318A15.2
RP11−305L7.7
RP11−59C5.3
LINC01588
RP5−1068E13.7 *
RP11−428O18.6
RP11−685G9.4
RP11−328C8.4 *
RP11−182J1.18
AC139100.4 *
LINC00665 *
ADPGK−AS1
RP11−1C8.5 *
CTC−550B14.7
RP11−46C24.7
RP11−299J3.8
PDCD4−AS1
NALT1
RP11−303E16.2
RP11−72M17.1
CTD−2622I13.3
RP11−73E17.2
RP11−586D19.2
LINC01307
UBA6−AS1
SNHG3
CTA−276F8.1
RP11−381K20.2
MAPKAPK5−AS1
RP11−849H4.4
C3orf35
ALOX12−AS1
LINC01140
CTB−13F3.1
AC006160.5
LA16c−390H2.4
RP11−316M1.12
Latent
Uninf.
Active
C
Module 1
Module 9
D

### Figure S2

| Gene name | qRT-PCT Primer sequence |
|---|---|
| RP11-1C8.5-QF<br>RP11-1C8.5-QR | CAGCCTTATTCTGAGGACCATAC<br>GCCAAATTGCCTAACCTCTTTC |
| RP11-328C8.4-QF<br>RP11-328C8.4-QR | CCACTGGCTGTGAGTTCAATA<br>GCAGTTTCATCAACCCATCAATAC |
| AC139100.4-QF<br>AC139100.4-QR | CTGGAAAGGACTCGAAGACAAA<br>GCTGGGTTGAGAGATGTGATG |
| RP5-1068E13.7-QF<br>RP5-1068E13.7-QR | TTATGCTAGGTGGAGAGGTAGG<br>TGCGTGTCTCTAACGAGTTTC |
| LINC00665-QF<br>LINC00665-QR | CACAGCAAGCCCCTGGAT<br>CAGATACTCAAGATGGGTGGTG |
| RP11-255C15.3-QF<br>RP11-255C15.3-QR | GAGCCCTCATGAATGGGATTAG<br>TTGCAGACTGCTGACTTCTC |
| NRAV-QF<br>NRAV-QR | GCTGTCTGGAGAGATGAAGAAA<br>CATCCCAGCTCTGTCACTTT |
| CTD-3222D19.12-QF<br>CTD-3222D19.12-QR | GATGCCTGTAATCCCATCTACTT<br>CACGACTTTGGCTCACTGTA |
| C21orf91-OT1-QF<br>C21orf91-OT1-QR | AATGTTTGGATGGCACAAGTC<br>CTCCTATCCAGTTTCTTTGCTTTAC |
| NEAT1-QF<br>NEAT1-QR | CCAGTTTTCCGAGAACCAAA<br>ATGCTGATCTGCTGCGTATG |
| GAPDH-QF<br>GAPDH-QF | GACAAGCTTCCCGTTCTCAG<br>GAGTCAACGGATTTGGTCGT |
| U6-QF<br>U6-QR | CTCGCTTTGGCAGCACA<br>AACGCTTCACGAATTTGCGT |
| 18SrRNA-QF<br>18SrRNA-QR | GGCCCTGTAATTGGAATGAGTC<br>CCAAGATCCAACTACGAGCTT |

### Table S1
